# Synaptic MEMOIR: mapping individual synapses of neurons with protein barcodes

**DOI:** 10.1101/2025.11.25.690442

**Authors:** Carsten H. Tischbirek, Linjing Fang, Duncan M. Chadly, Saori Lobbia, Jose David Aguirre, Yodai Takei, Chun-Hao Chen, Paul W. Sternberg, Carlos Lois, Michael B. Elowitz, Long Cai

**Affiliations:** Division of Biology and Biological Engineering, California Institute of Technology, Pasadena, CA, USA; Department of Biomedicine and University Hospital of Geriatric Medicine Felix Platter, University of Basel, Basel, Switzerland; Institute of Molecular and Cellular Biology, National Taiwan University, Taipei; Howard Hughes Medical Institute, California Institute of Technology, Pasadena, CA, USA

## Abstract

Obtaining wiring diagrams of brains has been a major achievement for neuroscience. However, an underlying challenge in connectomics is the fundamental tradeoff between the imaging resolution needed to resolve synapses and the volume of the brain that can be imaged. For example, electron microscopy (EM) visualizes synaptic sites with ~5 nm resolution, but is difficult to scale beyond volumes of 1 mm^3^. Here, we present Synaptic MEMOIR (Memory with Engineered Mutagenesis with Optical in situ Readout) that enables imaging of neuronal projections in animal brains with single-synapse resolution. Synaptic MEMOIR is built around three key design features. First, protein barcodes are transported to synapses to allow matching of synaptic barcodes to cell body barcodes without high resolution imaging and the error-prone process of tracing neuronal processes across long distances. Second, Synaptic MEMOIR uses continuous mutagenesis to generate a large diversity of barcodes to uniquely label cell bodies and synapses. Last, the timing of recombination and transport can be tuned to record synaptic age or projection information. Combining these features, we demonstrated projection mapping of 113 neurons in the *Drosophila melanogaster* optic lobe in a volume of 9.5 million μm^3^. Because synapses are identified by transported barcodes with optical microscopy at 300 nm resolution, this approach can potentially scale to much larger volumes, similar at least to those imaged in recent mouse brain transcriptomics atlases. In addition, synaptic MEMOIR can match barcodes across brain sections, and does not require tissue clearing to track long-range projections. Together, these results provide the foundation for a scalable optical image-based system for reconstructing the neural wiring diagrams of brains across different developmental stages, genetic backgrounds and perturbations.

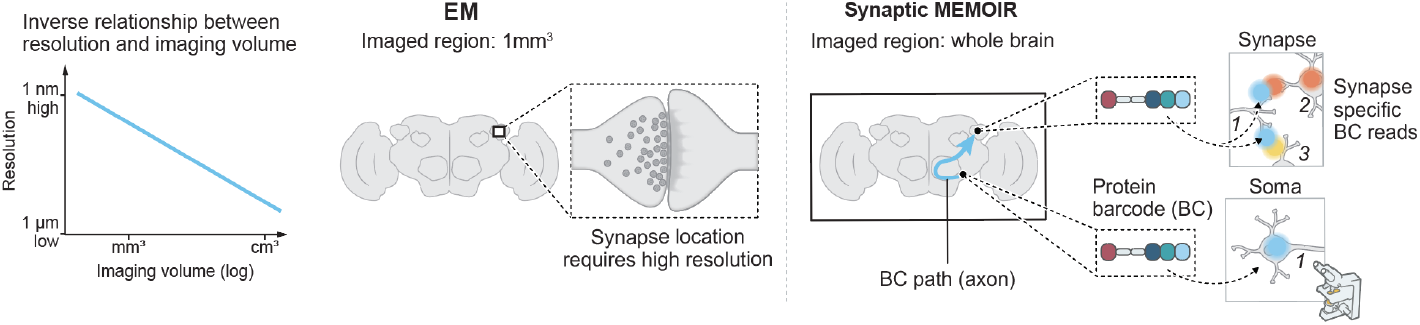

## Introduction

Mapping connections at the single-synapse level in brain circuits is essential for revealing circuit motifs that underlie brain computations. Electron microscopy (EM) has revolutionized connectomics research (Lichtman and Denk, 2011), showing success in mapping entire invertebrate brains and small brain volumes (~1 mm^3^) in mice (White et al., 1986; Zheng et al., 2018; Winding et al., 2023; Bae et al., 2025). Recent advances in optical imaging-based connectomics (Park et al., 2025; Tavakoli et al., 2025) have demonstrated alternative approaches using expansion microscopy and machine learning to segment cells and identify synapses in volumes up to 0.01 mm^3^.

However, the inverse relationship between imaging resolution and volume that can be imaged **(Fig. 1a)** limits scalability for mapping wiring diagrams in the brain. Identifying synapses currently requires high resolution imaging with EM (Dorkenwald et al., 2024; Bae et al., 2025) or expansion microscopy (Chen et al., 2015; Lillvis et al., 2022). At the same time, as resolution increases, the volume that can be imaged decreases proportionally (Lichtman and Denk, 2011; Lillvis et al., 2022). In EM with ~5 nm resolution, volumes of 1 mm^3^ were reconstructed, building upon more than a decade of heroic efforts (Kasthuri et al., 2015; Xu et al., 2017; Motta et al., 2019; Dorkenwald et al., 2024; Lin et al., 2024; Matsliah et al., 2024; Schlegel et al., 2024; Shapson-Coe et al., 2024; Bae et al., 2025). In recent expansion-based optical imaging experiments with ~50 nm resolution, 0.01 mm^3^ have been demonstrated (Lillvis et al., 2022; Tavakoli et al., 2025). Complete wiring diagrams of larger brains, as well as wiring diagrams of flies in different developmental stages and genetic backgrounds, require scaling up by 100- to 1000-fold in throughput.

**Figure 1.**
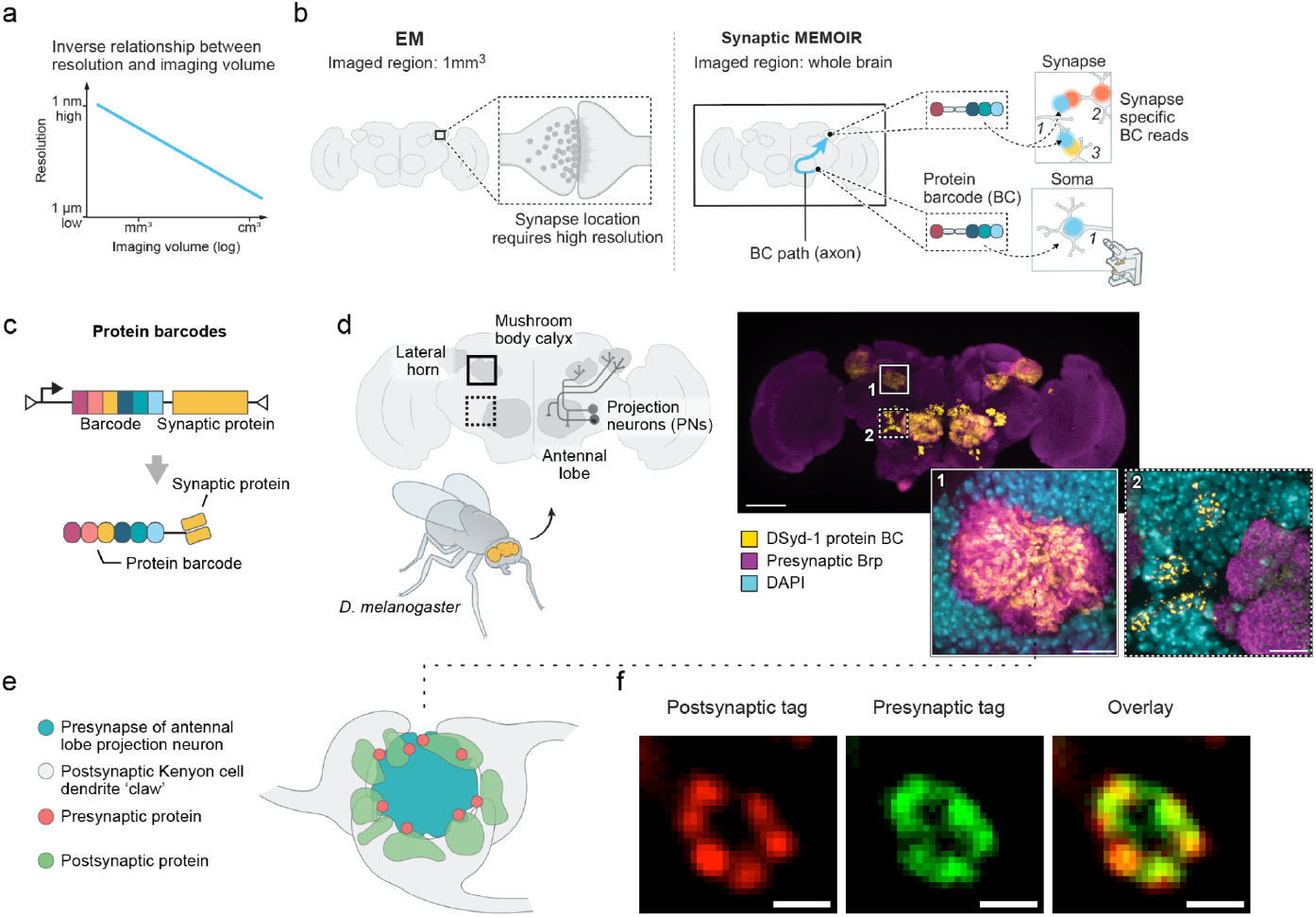
Overview of Synaptic MEMOIR. a, Conceptual illustration of the trade-off between imaging resolution and volume. b, Synaptic MEMOIR labels each neuron with a unique, genetically encoded barcode ID that is read out both at cell bodies and synapses, enabling the reconstruction of neuronal projection patterns with single-synapse resolution across long distances in brain tissue. c, Protein-barcode system design. A protein barcode cassette is fused and translated together with a synaptic localization protein. d, Validation of protein tag transport and synaptic localization in *D. melanogaster*. Schematic of antennal lobe projection neurons pathway with presynapses formed in the mushroom body calyx and the lateral horn. Maximum-intensity projection of a whole-mount brain labeled with anti-Brp (magenta), synaptic protein barcode tags (yellow) and DAPI (cyan). Scale bar = 100 um. Insets show protein tag signals detected in mushroom body calyx synapses (1, white box in overview image) and antennal lobe projection neuron cell bodies (2, white dotted line box in overview image). Scale bars = 10 μm. e, Schematic of a calyx synapse indicating pre- and postsynaptic compartments. f, Example co-localization of presynaptic and postsynaptic protein-barcode fusion tags using DSyd-1 and Drep-2, respectively. Scale bar = 1 μm.

Here, we propose a new approach to overcome this volume-resolution trade-off called Synaptic MEMOIR (Memory with Engineered Mutagenesis with Optical in situ Readout) (**Fig. 1b**). Building upon MEMOIR tools we previously developed to continuously mutagenize genetic barcodes for lineage recording (Frieda et al., 2017; Chow et al., 2021), we engineered cells to express unique protein barcodes that can be identified in the cell bodies and transported to the synaptic sites of neurons. In this fashion, the synaptic barcodes can be matched to the cell body barcodes to uniquely identify the synapses to their corresponding cell body. In principle, with a large diversity of barcodes, each neuron in the brain can be uniquely labeled, allowing scalable identification of synaptic projection and connectivity across large brain regions.

The key design feature of Synaptic MEMOIR is that transported barcodes **(Fig. 1b)** enable identification of synapses with diffraction-limited optical imaging (~300 nm resolution). At this lower resolution compared to EM, 1000 times larger imaging volumes (~1 cm^3^) become possible. For comparison, mouse brain transcriptomics atlas efforts (Zhang et al., 2023; Yao et al., 2023; Shi et al., 2023) used similar optical sequential imaging methods to image entire mouse brains of approximately 1 cm^3^. The density of synapses across the mouse brain is estimated to be 1 synapse per µm^3^ (Santuy et al., 2020), which indicates that if transported barcodes are used, lower resolution imaging methods can distinguish different synapses. Density can be reduced further by sequential imaging, similar to our efforts to image transcriptome-scale gene expression (Eng et al., 2019; Takei et al., 2021, 2025), especially when barcodes are diverse.

In addition, removing the need to trace neuronal processes eliminates an error-prone and labor-intensive step that contributes to the limitation of current connectomics approaches to 1 mm^3^ volumes.

Beyond transported barcodes, our design has two additional key features to allow synapse mapping to be scalable. First, barcodes must reach synapses efficiently and uniformly. RNA barcodes have been used in pioneering methods such as MAPseq and BARseq (Chen et al., 2019; Kebschull et al., 2016) to map projections in the mouse brain, but have important limitations. In particular, the sparseness of RNA expression limits the resolution of the projection analysis to axons and brain regions and cannot readily resolve individual synapses. By contrast, protein levels are typically approximately a thousand-fold higher than RNA levels (Schwanhäusser et al., 2011), which increases the likelihood of uniform synaptic labeling within a neuron. Considering that neurons can form thousands to hundreds of thousands of synapses, it is unlikely that RNA barcodes can be expressed at levels sufficient to be uniformly distributed across this many synapses, making a protein barcoding system highly advantageous. Protein barcodes also simplify the genetic implementation of the system. Barcode fusions with synaptic proteins remove the need for a separate transporter system, and combining multiple barcode elements in a single protein avoids providing each element with its own promoter and transporter, which reduces the overall sequence size and burden. The disadvantage of protein detection is that it requires antibodies, which are less robust than oligonucleotide probes used for RNA barcode detection. However, this drawback is less consequential than the risk of inadequate expression with RNA barcodes.

As a second criteria, barcodes must provide sufficient diversity to uniquely label neurons. To label most neurons across the brain (100,000 neurons in the fly and 70 million in the mouse brain), we need barcodes with 100- to 1000-fold larger diversity to minimize the chances of two neurons being labeled by the same barcodes. With protein barcodes, only around 33 epitopes are needed to achieve this diversity (2^33^ ≈ 10^10^). In Synaptic MEMOIR, we used recombinases to create diversity through continuous mutagenesis of a single genomically integrated barcode site. The integration of a barcode array, consisting of multiple epitope tags translated as a single protein, ensures that neurons can express the synaptic barcodes as a whole and that the synapses are filled uniformly. We further showed that this approach is scalable with multiple barcode arrays by demonstrating that two distinct genomic integrated barcodes are both expressed and present at individual synapses.

In proof-of-principle experiments, we show that Synaptic MEMOIR barcodes can be edited, expressed, transported and detected in the *Drosophila* optic lobe. *Drosophila* was chosen because it provided well-defined neuronal connectivity and a readily available toolset for genetic manipulation, and allowed for ease of maintenance during the pandemic. We envision Synaptic MEMOIR as the foundation for a highly scalable system to perform synapse-resolved neuronal projection mapping compatible with any organism with suitable genetic tools.

## Results

### 1. Active transport of protein barcodes to synaptic sites

For a transported synapse mapping system to be scalable, barcodes must reach synaptic terminals efficiently and uniformly. We expressed a protein barcode, consisting of multiple epitope tags, fused to a presynaptic protein (DSyd-1) in *Drosophila melanogaster* antennal lobe projection neurons using the GH146-GAL4 driver line **(Fig. 1c)**. We selected the presynaptic protein DSyd-1 (Owald et al., 2010) as a transporter, as it localized efficiently to active zones in screening experiments that included different GFP-fusion synaptic proteins such as Rab-3 and Bruchpilot (Brp). Localization of DSyd-1 fused barcode, detected by antibody labeling, was examined by co-labeling with antibodies against Brp, a marker of presynaptic sites, and with DAPI to mark nuclei surrounding the antennal lobe glomeruli **(Fig. 1d)**.

As expected, a strong barcode signal was detected in the cell bodies of the antennal lobe projection neurons surrounding the antennal lobe. Barcode signal was also observed in the antennal lobe glomerular regions, the calyx, and the lateral horn-structures containing projection neuron presynaptic terminals, suggesting that barcodes were being consistently trafficked to distant synaptic terminals. Colocalization analysis showed that almost all barcode puncta overlapped with Brp-positive synaptic sites **(Fig. S1)**, which indicates specific transport to presynaptic compartments rather than passive diffusion. Axons had significantly weaker signals. This “clean” trafficking to synapses supports the use of protein barcodes for mapping synaptic sites far from the soma.

Both presynaptic and postsynaptic terminals can be labeled by transported barcodes (**Fig. 1e, f**) and matched to identify synapses. In the *Drosophila* mushroom body, synapses have a distinct morphology **(Fig. 1e)**. Staining of the postsynaptic transport tags fused to Drep-2 and a presynaptic tag to DSyd-1 protein show this clear morphology, indicating that the pre- and post-synapse can be accurately identified using this approach **(Fig. 1f)**.

### 2. Epitope selection for protein barcodes

To implement synaptic MEMOIR, we had to identify multiple epitope tags as orthogonally detectable components of protein barcodes. We screened a library of epitope tags that are also orthogonal to endogenous proteins. We curated a list of 50 epitopes, such as Myc- and HA-tags, that can be targeted by commercially available antibodies. We generated single epitope tag cell lines for each epitope and tested the antibodies.

To multiplex the readout of barcode elements *in situ*, we adapted an oligo-conjugated antibody detection strategy compatible with sequential fluorescence hybridization **(Fig. S2)**. Each epitope tag was recognized by a tag-specific antibody covalently linked to a single-stranded DNA oligonucleotide with a 15 nt sequence unique to that tag. Antibodies were conjugated via amine-reactive linkers to amine-modified oligos using established protocols (Black et al., 2021; Takei et al., 2021). From the 50 epitope tags screened, we obtained 12 tags that provided good signals with oligo-conjugated antibodies.

### 3. Recombination of barcodes generates diversity

To generate a different barcode in each neuron, we constructed genetically-encoded protein epitope arrays that undergo stochastic, site-specific deletion of constituent elements mediated by the serine integrase Bxb1 (Chow et al., 2021). We expressed the synaptically transported barcode array from a single genomic integration site to simplify delivery, ensure stoichiometric production of all barcode elements (epitopes), and reduce variability of trafficking to synaptic sites **(Fig. 2a)**.

**Figure 2.**
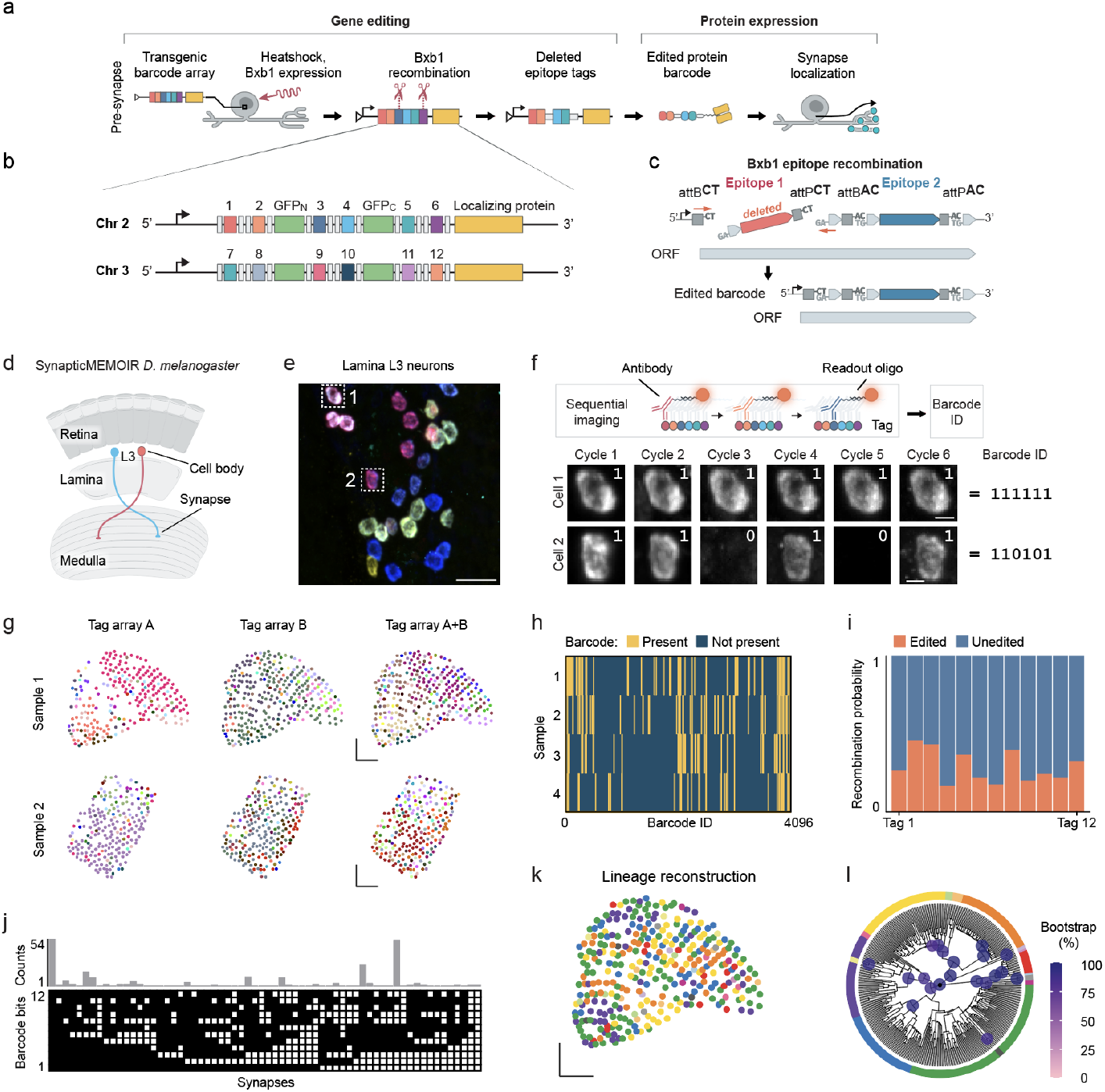
Stochastic recombination generates diverse protein barcodes that can be detected *in situ*. a, Experimental overview. For stochastic gene editing, a Bxb1 integrase enzyme performs stochastic barcode DNA recombination. Barcodes are then expressed and trafficked to their target region. b, Synaptic barcodes consist of a presynaptic protein DSyd-1, a GFP backbone and six unique epitope tags each flanked by orthogonal Bxb1 editing sites. c, Overview of the molecular processes involved in protein tag expression. d, Schematic of the *Drosophila* optic lobe. e, L3 neuron cell bodies after recombination shown here as a color-coded overlay image of six imaging rounds. Scale bar = 10 μm. f, Sequential imaging allows multiplex detection of protein barcode bits *in situ*. Magnified view of regions 1 and 2 marked in panel (e) showing individual imaging cycles with binary barcode readout with signal = 1, no signal = 0. Scale bar = 2 μm. g, Barcode masks of L3 neuron projections in the optic lobe. Colors represent individual barcode IDs. Tag A and B are two orthogonal barcode arrays with six epitope tags each, with more barcode diversity observed in the combined dataset. Scale bar = 25 μm. h, Summary of detected barcode IDs across four optic lobes with all possible variants of the 12-bit (4096) barcode on the x-axis. i, Per-tag recombination percentages obtained from four optic lobe experiments. j, Barcode diversity and frequency from sample 1 of (h). Top: histogram of barcode repeat counts. Bottom: binary barcode matrix (black, unedited; white, edited). k, Synaptic MEMOIR reveals lineage information of synaptic projections, shown here color-coded according to barcode lineage. Colors represent clones supported by at least 80% transfer score as computed from posterior samples generated by the Bayesian Evolutionary Analysis by Sampling Trees platform (BEAST2). Scale bar = 25 μm. l, Circular lineage tree diagram of the sample shown in panel (k). Color code used in panel (k) is shown as the outer ring of the circular lineage tree plot. Colored circles on interior nodes of the tree represent the phylogenetic transfer score, shown for clades with at least 80% support.

Barcode diversity was generated by Bxb1 recombination at *att* sites distinguished by central dinucleotide variants (Stark et al., 1992; Ghosh et al., 2003; Chow et al., 2021), which control pairing and orientation of the recombination. Of the 16 theoretical dinucleotide pairs, we implemented six orthogonal pairs to avoid cross-recombination **(Fig. S3)**. We excluded pairs that (i) exhibited cross-reactivity enabling non-cognate recombination, or (ii) were palindromic and prone to inversions besides deletions. Also, spacer nucleotides in the design ensured that recombination events of the attB site and attP site did not cause frame shifts leading to nonsensical translation and premature stop codons. The attB and attP sequences themselves are in frame so that they do not code for a stop codon prematurely disrupting Synaptic MEMOIR barcode translation **(Fig. S3)**. This filtering produced a library in which recombination predominantly resolved as clean deletions of individual elements.

We integrated two barcode arrays consisting of 6 epitope tags each into chromosome II and III of *Drosophila*, using the 12 orthogonal epitope tags from the screen **(Fig. 2b)**. The dual-tag configuration has a theoretical diversity to 4096 possible barcodes per cell. Each barcode element comprised tandem repeats of short, linear epitope tags **(Table S1)**. Elements were embedded within a GFP scaffold (Viswanathan et al., 2015) to aid visualization in functional assays. Specifically, tag arrays were positioned at the GFP N-terminus, within an internal loop, and at the C-terminus, allowing modular targeting via appended localization signals. Recombination-induced deletions were designed to be frame-safe and generated attL/attR junctions whose residual “scars” preserved the reading frame and avoided stop codons **(Fig. 2c, Fig. S4)**, maintaining transcriptional and translational competence of the barcode protein after element loss. Thus, the encoded protein remains intact while the presence and absence pattern of tag elements records the recombination history.

### 4. In situ detection of barcode diversity and lineage tracing in D. melanogaster neurons

We next induced Bxb1-mediated recombination to generate barcode diversity in the *Drosophila* optic lobe. We expressed the barcodes in L3-neurons using a R14B07-GAL4 line **(Fig. 2d)**, with the Bxb1 recombinase placed under control of a heat shock promoter. In this configuration, all three components - the GAL4 driver, the barcode protein, and the heat shock-inducible Bxb1 recombinase (Chow et al., 2021) - were combined in the same genetic background. Larvae were heat-shocked 48 hours after egg laying, and barcode patterns were analyzed in adult flies. We first PCR-amplified the barcode loci from brain sections to show that recombination occurred randomly, obtaining a ladder of edited barcode states **(Fig. S5)**.

To image the recombined barcodes *in situ*, the samples (**Fig. 2e**) were co-labeled with the full set of tag-specific oligo-conjugated antibodies corresponding to the barcode’s epitope repertoire **(Fig. 2f)**. Sequential readout proceeded as follows: in each cycle, a fluorophore-labeled, complementary 15 nt “readout probe” was hybridized to a single antibody-conjugated oligo. Following imaging, the readout probe was stripped with a brief formamide wash analogous to probe removal steps in seqFISH (Shah et al., 2018; Eng et al., 2019) and the next cycle was initiated with a readout probe complementary to a different antibody-conjugated oligo. This process was repeated until all barcode elements were imaged. This approach enables orthogonal, cycle-by-cycle binary detection of the presence (=1) or absence (=0) of multiple protein barcode elements within the same cell to generate a binary barcode for the cell body or synapse **(Fig. 2f)**, without spectral crowding or cross-talk between readout channels. The strategy leverages the high protein abundance achievable from single-copy monogenic barcodes, making them detectable in small synaptic sites where RNA barcodes may be unstable or below the detection threshold.

Confocal imaging of four optic lobes revealed stochastic recombination events, with unique barcode combinations observed in the synaptic terminals of lamina neurons in each sample (**Fig. 2g-h**). Individual epitope tag elements recombined with similar efficiencies, and the overall fraction of recombined barcode elements per neuron averaged approximately 27% **(Fig. 2i)**. Recombination levels were tunable: varying heat shock intensity modulated the extent of barcode element deletion, enabling controlled diversity generation **(Fig. S6)**.

Importantly, the two barcode arrays were not only recombined independently from each other **(Fig. 2j**) but were transported to the same synapses. Both tag arrays were observed at 89% of synapses **(Fig. 2g)**. Similar barcode diversity was observed for both tandem tags **(Fig. 2h)**. Tag array A had 49 out of a total of 64 combinations, and tag array B had 51 out of 64 combinations. The two tag arrays together provided 216 unique barcodes out of 4096 unique possible combinations **(Fig. 2h)** in an example dataset of 884 synapses, with duplicate barcodes reflecting the limited barcoding space. An additional experiment using recombination of a 11-tag system also showed independent editing of each epitope tag **(Fig. S7)**. This indicates that increasing the barcode capacity with 33 tags can be achieved by introducing separate genomically integrated barcode sites, each containing 11 epitope tags.

An intrinsic advantage of the genetically-encoded protein-based barcodes lies in their heritability. Once recombined, the barcode state is stably inherited by daughter neurons, enabling simultaneous lineage inference and projection mapping. We analyzed barcode patterns at synaptic sites in *Drosophila* L3 lamina neurons using the dual tag array recombination system (**Fig. 2k-l**). The resulting lineage trees revealed groups of L3 neuron synapses with identical or near identical barcodes **(Fig. 2k-l)**. Spatial mapping of the related L3 neurons showed distinct columns, sometimes adjacent but frequently separated across the medulla **(Fig. 2k)**. These results confirm the feasibility of combining lineage and projection readout at distal synaptic sites.

### 5. Accumulation of synaptic barcodes can measure synaptic age

We next set out to demonstrate that synaptic barcodes and cell body barcodes can be matched to generate synaptic projection information. We used a GH146-GAL4 driver line to express MEMOIR barcodes in antennal lobe projection neurons (**Fig. 3a**) and used heat shock induction to express Bxb1 integrase for barcode recombination (Chow et al., 2021). To our disappointment, we observed little matching between the synaptic and cell body barcodes (about 11.6% of synapses share a barcode with a cell body).

**Figure 3.**
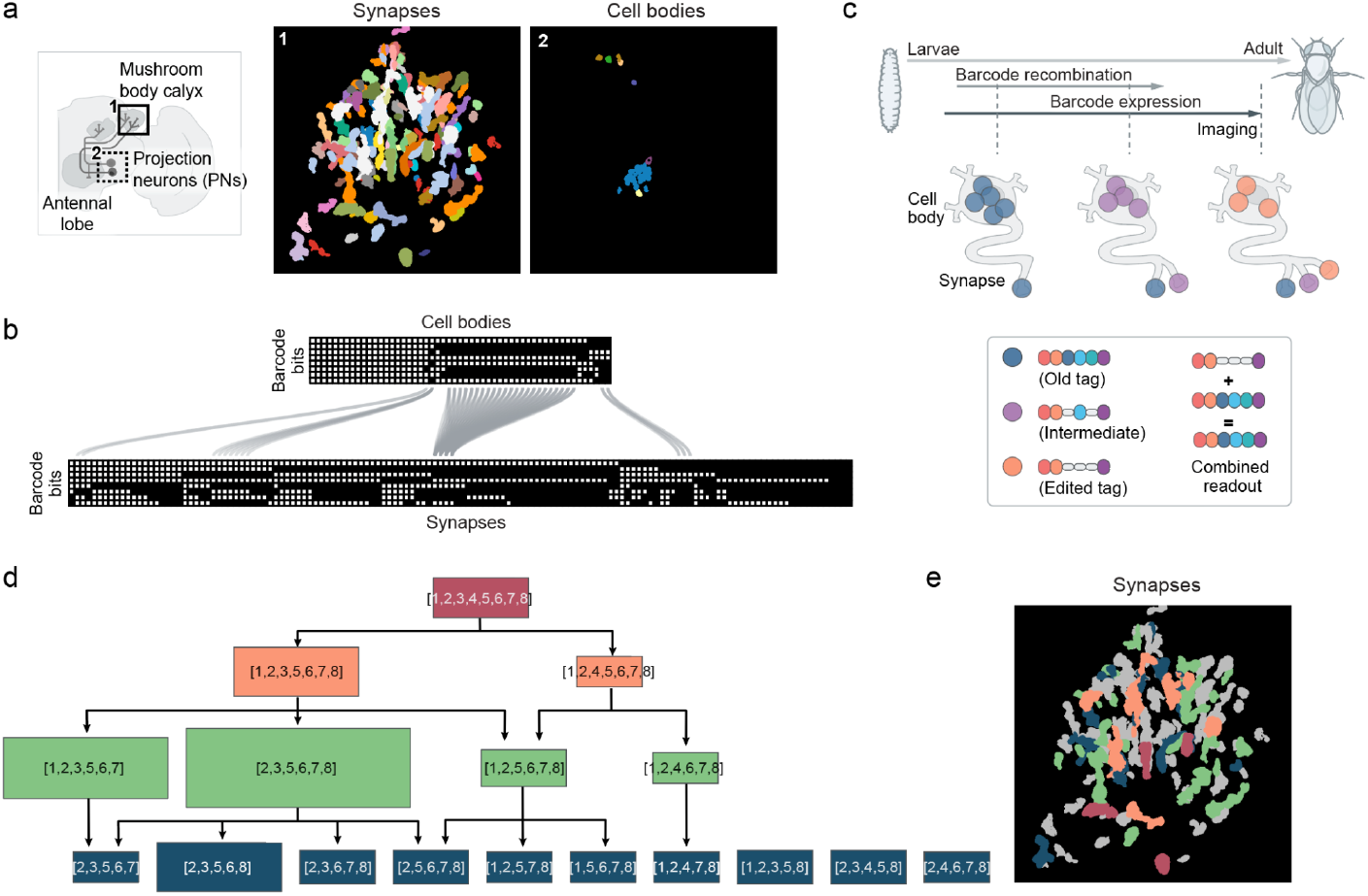
Accumulation of protein barcode in the synapse can reflect synaptic age. a, Segmented synapses and cell bodies of antennal lobe projection neurons images at the regions indicated in the schematic on the left, Synapses in the left image and cell bodies in the right image were color coded according to their barcode ID. b, Barcode bits of cell bodies identified near the antennal lobes connected with calyx synapses when a barcode is identical in cell bodies and synapses. Note the low amount of barcode matching between both compartments. c, Simple accumulation model illustrating how existing, unedited barcodes which are positive for all tags can dominate the detectable signal and obscure subsequently edited barcode states. d, Lineage tree of “old” Synaptic MEMOIR barcodes that are related to each other in the sample. Numbers in boxes correspond to epitopes (or image cycles) that had a positive barcode signal for the synapse. Boxes are scaled according to the number of synapses sharing the barcode in the sample. e, Segmented synapses in the Calyx color-coded according to the lineage tree shown in panel (d). Grey objects correspond to regions with no significant relationship to the lineage tree

We hypothesized that the discrepancy between the soma and synaptic barcodes could be due to the accumulation of “old” barcodes in the synapse even after the genomic copy of the barcodes has been further edited. This problem occurs when recombination and transport of the barcodes are co-occurring across a period of multiple days during heat shock and expression of the integrase and barcodes. Consistent with this hypothesis, cell body barcodes appeared to be more edited compared to the synaptic barcodes (70.3% (53 cells) vs 46.7% (138 synapses) for the whole-mount brain shown in **Fig. 3b**). In addition, observations of completely unedited barcodes at the synapse, even though all cell body barcodes contained some form of edits, further suggest that the protein degradation rate at the synapse is extremely slow, i.e. once a protein barcode is transported to the synapse, it is likely to remain.

Thus, any subset of synaptic barcodes that can match to a cell barcode could be due to accumulation of barcodes as the cell progressed through different stages of editing. For example, synaptic barcodes with only epitope 8 remaining cannot possibly arise from cells that still had epitope 4 (or any other epitopes besides 8) still present. Those synapses may have formed from a neuron that had only epitope 8 unedited at one stage on its way to a completely edited cell barcode.

Interestingly, the accumulation of distinct synaptic barcodes can provide information on the “age” of synapses **(Fig. 3c)**. Completely unedited barcodes were present at several synapses, suggesting these are the oldest synapses. These synapses with unedited barcodes likely correspond to projections to the calyx that were formed prior to heat shock induction of integrases. We then constructed a tree of edits to the synaptic barcodes as new synapses form **(Fig. 3d)**. Instead of observing all possible edits to the initial unedited barcode, we detected only deletion of epitope 3 and 4 in synapses (top row of **Fig. 3d**). Subsequent edits also occur for a subset of the epitopes instead of all possible edits, suggesting progressive editing in neurons that are projected to the calyx as the brain develops.

### 6. Matching of synaptic and cell body barcodes

To improve the design to generate more matching between somatic and synaptic barcodes, we implemented a two-phase strategy that confines barcode expression to a defined temporal window after heat shock induction of Bxb1 recombinations. This phased strategy reduced the accumulation of barcodes at different stages of recombination in the synapse and improved matching to the soma barcodes **(Fig. 4a)**. In the first editing phase, the Bxb1 integrase was transiently induced from a heat shock promoter to stochastically recombine the genomic barcode cassettes. In the subsequent expression phase, barcode proteins were expressed under the control of the *Drosophila* Auxin-inducible Gene Expression System (AGES) (McClure et al., 2022), which enables auxin-dependent activation of GAL4-driven transcription. The samples were heatshocked multiple times to increase recombination of the barcodes with an overall lower count of the cells. As expected, following the increased number of heat shocks, we saw a much sparser expression of barcodes compared to other heat shock conditions (**Fig. S8**).

**Figure 4.**
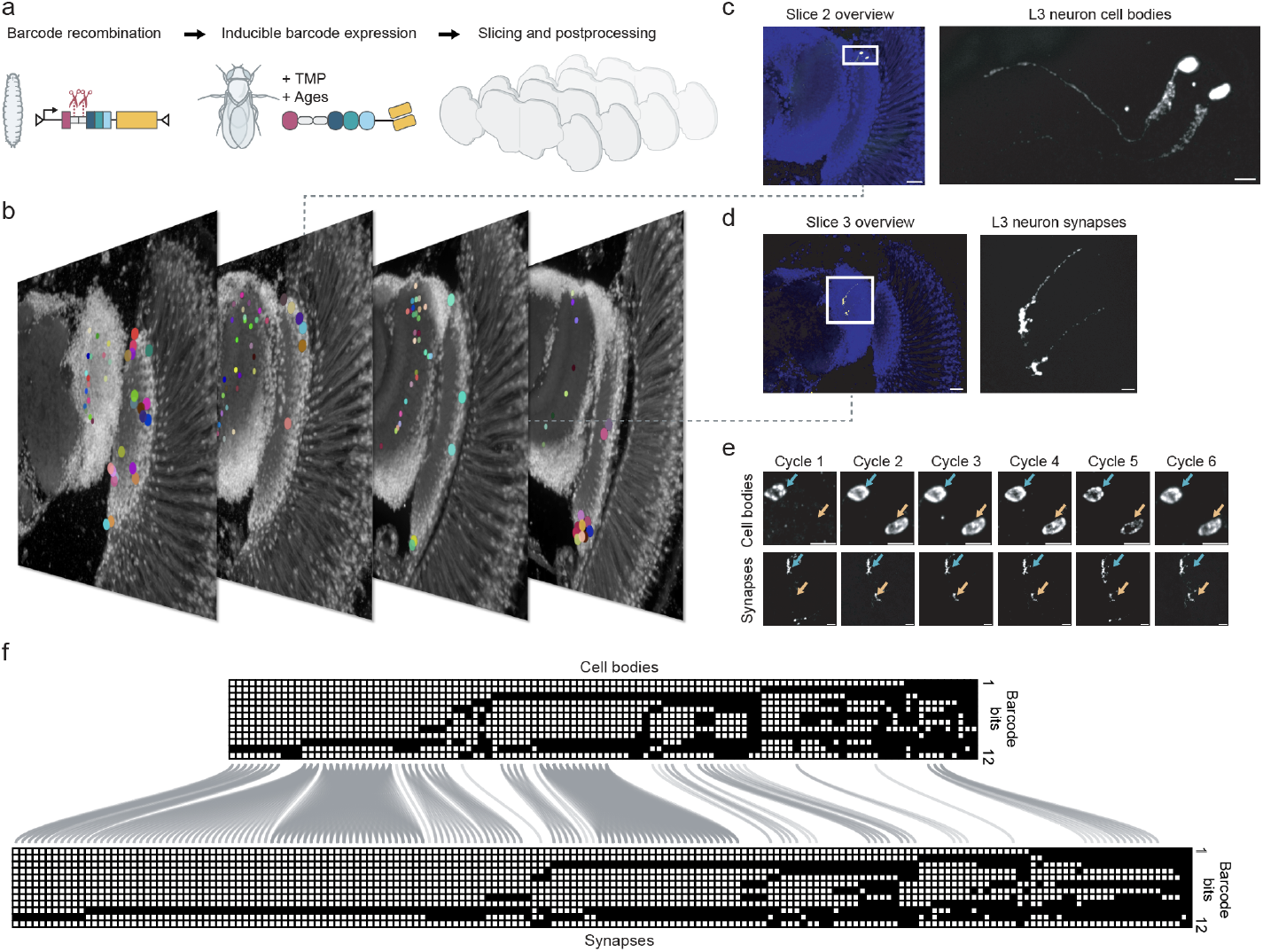
Mapping neuronal projections across discontinuous sections with Synaptic MEMOIR. a, Two-phase inducible Synaptic MEMOIR system workflow. Barcodes are expressed under GAL4/UAS with temporal control via the TMP-stabilized degron and AGES systems, enabling barcode induction after barcode recombination. b, Overview of segmented cell bodies (large dots) and synapses (small dots) across serial, discontinuous sections (~25 μm thick) of an optic lobe with sparse barcode tag expression following multiple heat shock induction of Bxb1. c, Cells in one z-section of slice 2 from (b), shown for one imaging cycle. Magnified region with barcode-positive cell bodies with processes faintly labeled. Scale bars 20 μm (overview) and 5 μm (magnified). d, Synapses corresponding to cells in (c) imaged in slice 3, box region is magnified. e, Sequential imaging cycles for the cells and synapses in (c-d), showing distinct barcode states across cycles. Note absence of signal in the cell bodies and synapses of the lower cell in corresponding first imaging cycle as marked by yellow arrows. Scale bar 2 μm. f, Binary barcode matrices for cell bodies and synapses across slices, with lines indicating that 75.9% of cell bodies and synapses have matching barcodes.

With this system in the optic lobe **(Fig. 4b)**, we observed significant matching between the cell body and synapse barcodes. Across all 4 physical slices spanning ~9.5 million μm^3^ of the fly brain, we observed 113 cell bodies and 166 synapses. 75.9% of the synaptic barcodes have corresponding cell body barcodes. This is consistent with our sectioning and imaging of approximately three-quarters of the optic lobe. Most of the matches are between groups of synapses and groups of cell bodies carrying the same barcodes. Single synapses and single cell bodies matched in 10 instances, due to the small number of epitope tags and barcoding space available.

We validated the barcode matching by tracing the neuronal processes, which showed a faint signal in one of the epitope channels. We can trace the processes across two physical sections of the *Drosophila* brain to connect the synapses with the cell bodies. In one field of view **(Fig. 4c)**, we observed two cell bodies with distinct barcodes **(Fig. 4e**, in the lower cell one of the epitopes was deleted by Bxb1 recombination) and, correspondingly, in another section two synapses of the sparsely labeled sample had the same set of barcodes (**Fig. 4d**). Another example is shown for a cell that was traceable within a single section (**Fig. S9**), validating the projection matching of the Synaptic MEMOIR barcodes.

Matching across sample sections of the barcodes suggest that it is no longer necessary to use whole-mount brains and clearing methods to trace the neuronal processes from the cell bodies to the synapse. Additional epitope tags beyond the currently identified 12 tags would dramatically increase the barcode diversity and allow many more unique matches between individual neurons and their synapses.

## Discussion

Synaptic MEMOIR demonstrates that (i) protein barcodes can be transported to synapses, (ii) a diverse set of barcodes can be generated by the MEMOIR system from a single genomically integrated site, and (iii) the timing of recombination, expression, and transport can be controlled to match synaptic barcodes with cell body barcodes. This new experimental paradigm overcomes the resolution-volume tradeoff to enable scalable imaging based mapping of individual synapses to their neurons. We mapped 113 cell bodies in ~0.01 mm^3^ volume, comparable in number of neurons and total volume to recent work (Park et al., 2025; Tavakoli et al., 2025) in optical connectomics, albeit with fewer synapses in a sparser sample.

Synaptic MEMOIR has the potential to scale to significantly larger numbers of cells and imaging volumes. The limitation of the current method is mainly due to the small number of epitopes (12) incorporated in the barcodes (2^12^ = 4096). By screening and identifying additional epitope tags, a much larger number of neurons can be uniquely labeled, minimizing the sharing of the same barcodes in multiple cells encountered in the present work. For example, 33 epitope tags can reach a maximum diversity of 10^10^ barcodes, making tagging entire fly (10^5^), zebrafish (10^6^-10^7^) and mouse brains (10^8^) possible. In addition, we have only demonstrated presynaptic site matching with cell bodies because of the limited number of epitopes. Adding postsynaptic barcodes would allow connectivity between cells to be matched, but would require an additional 33 epitopes. Importantly, especially for connectivity and projection mapping in high density samples, a larger barcoding space results in sparser sequential images, which “dilutes” the signal density, similar to the effect demonstrated in seqFISH+ (Eng et al., 2019; Takei et al., 2021, 2025).

Synaptic MEMOIR complements existing approaches to map neuronal processes and projections. Process-filling approaches such as Brainbow or Bitbow (Livet et al., 2007; Li et al., 2021) can provide distinct labeling of many neurons and reveal clonal neuronal populations. Similarly, recent work also explored protein-based cell-filling with antibody readout and expansion microscopy (Park et al., 2025). These methods do not solve the scalability challenge because they still require super-resolution imaging to identify synapses. In addition, long-range tracing across large volumes, even with unique identifiers, requires extensive image registration and manual or computational tracing. Methods using pre-formed RNA barcodes expressed from viral libraries, such as BARseq and MAPseq (Chen et al., 2019; Kebschull et al., 2016) can map long-range projections to cell body locations, but did not provide single-synapse resolution. Synaptic MEMOIR complements these approaches by reading out dynamically generated barcodes at individual synapses and matching them to the cell bodies of the same neurons to overcome the scalability problem.

The ability to match barcodes at sites distant from the nucleus removes the need for tissue clearing methods to trace long-range projections and avoids the labor-intensive alignment of individual slices. We demonstrated that synapses can be matched to cell bodies in different physical sections of the brain (**Fig. 4b**). Thus, the entire projection and connectivity patterns of the brain could be reconstructed using serial sections that are imaged separately from each other, rather than in whole-mount cleared brains.

The significant advantage of working with sections is the compatibility with recently developed spatial multi-omics experiments (Shah et al., 2018; Kishi et al., 2019; Takei et al., 2021; Gandin et al., 2025; Alon et al., 2021; Su et al., 2020; Takei et al., 2025) needed to characterize the molecular state of the cells along with lineage, projection and connectivity in the same cell. This opens possibilities for directly linking molecular identity to projection patterns and synaptic connectivity. In the nervous system, such an approach could be used to map the relationship between developmental lineage and wiring topology in a single experiment. Synaptic MEMOIR can also potentially integrate with calcium imaging (Grienberger and Konnerth, 2012; Chen et al., 2013) to determine firing patterns of neurons overlaid on their connectivity maps. Application of this integrated and scalable approach can be powerful to study brain wiring diagrams across different developmental stages, genetic backgrounds and perturbation conditions.

## Acknowledgments

We thank Inna-Marie Strazhnik for help with figures, Jina Yun, Sihui Yang and Ivan Luu for technical support, Aubrie De La Cruz for help with initial *Drosophila* work, James Linton and Kirsten Frieda for insightful discussions, the Caltech Proteome Exploration Laboratory for access to laboratory equipment, and Jenna Sternberg for help with text editing. We also thank members of Elowitz, Lois, Sternberg and Cai labs for helpful discussions and critical feedback. Stocks obtained from the Bloomington Drosophila Stock Center (NIH P40OD018537) were used in this study. This project was supported by a grant from the Allen Foundation Discovery Center and NIH TR01MH116508.

## Author Contributions

C.H.T. and L.C. conceived the idea and designed experiments. C.H.T., S.L., and J.D.A. prepared and validated experimental materials and designed probes with input from Y.T.. C.-H. C. prepared *C. elegans* experiments with input from P.W.S.. C.H.T., S.L., and J.D.A. performed experiments. L.F. and C.H.T. jointly analyzed the Synaptic MEMOIR data with input from L.C., P.W.S., C.L., and M.B.E.. D.M.C. and L.F. performed lineage analysis. C.H.T., L.F., D.M.C., and L.C. made the figures. C.H.T. and L.C. wrote the manuscript with input from all authors.

## References

Alon, S., Goodwin, D.R., Sinha, A., Wassie, A.T., Chen, F., Daugharthy, E.R., Bando, Y., Kajita, A., Xue, A.G., Marrett, K., et al. (2021). Expansion sequencing: Spatially precise in situ transcriptomics in intact biological systems. Science 371, eaax2656. 10.1126/science.aax2656.

Bae, J.A., Baptiste, M., Baptiste, M.R., Bishop, C.A., Bodor, A.L., Brittain, D., Brooks, V., Buchanan, J., Bumbarger, D.J., Castro, M.A., et al. (2025). Functional connectomics spanning multiple areas of mouse visual cortex. Nature 640, 435–447. 10.1038/s41586-025-08790-w.

Black, S., Phillips, D., Hickey, J.W., Kennedy-Darling, J., Venkataraaman, V.G., Samusik, N., Goltsev, Y., Schürch, C.M., and Nolan, G.P. (2021). CODEX multiplexed tissue imaging with DNA-conjugated antibodies. Nat. Protoc. 16, 3802–3835. 10.1038/s41596-021-00556-8.

Chen, F., Tillberg, P.W., and Boyden, E.S. (2015). Expansion microscopy. Science 347, 543–548. 10.1126/science.1260088.

Chen, T.-W., Wardill, T.J., Sun, Y., Pulver, S.R., Renninger, S.L., Baohan, A., Schreiter, E.R., Kerr, R.A., Orger, M.B., Jayaraman, V., et al. (2013). Ultrasensitive fluorescent proteins for imaging neuronal activity. Nature 499, 295–300. 10.1038/nature12354.

Chen, X., Sun, Y.-C., Zhan, H., Kebschull, J.M., Fischer, S., Matho, K., Huang, Z.J., Gillis, J., and Zador, A.M. (2019). High-Throughput Mapping of Long-Range Neuronal Projection Using In Situ Sequencing. Cell 179, 772–786.e19. 10.1016/j.cell.2019.09.023.

Chow, K.-H.K., Budde, M.W., Granados, A.A., Cabrera, M., Yoon, S., Cho, S., Huang, T., Koulena, N., Frieda, K.L., Cai, L., et al. (2021). Imaging cell lineage with a synthetic digital recording system. Science 372, eabb3099. 10.1126/science.abb3099.

Dorkenwald, S., Matsliah, A., Sterling, A.R., Schlegel, P., Yu, S., McKellar, C.E., Lin, A., Costa, M., Eichler, K., Yin, Y., et al. (2024). Neuronal wiring diagram of an adult brain. Nature 634, 124–138. 10.1038/s41586-024-07558-y.

Eng, C.-H.L., Lawson, M., Zhu, Q., Dries, R., Koulena, N., Takei, Y., Yun, J., Cronin, C., Karp, C., Yuan, G.-C., et al. (2019). Transcriptome-scale super-resolved imaging in tissues by RNA seqFISH. Nature 568, 235–239. 10.1038/s41586-019-1049-y.

Frieda, K.L., Linton, J.M., Hormoz, S., Choi, J., Chow, K.-H.K., Singer, Z.S., Budde, M.W., Elowitz, M.B., and Cai, L. (2017). Synthetic recording and in situ readout of lineage information in single cells. Nature 541, 107–111. 10.1038/nature20777.

Gandin, V., Kim, J., Yang, L.-Z., Lian, Y., Kawase, T., Hu, A., Rokicki, K., Fleishman, G., Tillberg, P., Castrejon, A.A., et al. (2025). Deep-tissue transcriptomics and subcellular imaging at high spatial resolution. Science 388, eadq2084. 10.1126/science.adq2084.

Ghosh, P., Kim, A.I., and Hatfull, G.F. (2003). The Orientation of Mycobacteriophage Bxb1 Integration Is Solely Dependent on the Central Dinucleotide of attP and attB. Mol. Cell 12, 1101–1111. 10.1016/S1097-2765(03)00444-1.

Grienberger, C., and Konnerth, A. (2012). Imaging calcium in neurons. Neuron 73, 862–885. 10.1016/j.neuron.2012.02.011.

Kasthuri, N., Hayworth, K.J., Berger, D.R., Schalek, R.L., Conchello, J.A., Knowles-Barley, S., Lee, D., Vázquez-Reina, A., Kaynig, V., Jones, T.R., et al. (2015). Saturated Reconstruction of a Volume of Neocortex. Cell 162, 648–661. 10.1016/j.cell.2015.06.054.

Kebschull, J.M., Garcia da Silva, P., Reid, A.P., Peikon, I.D., Albeanu, D.F., and Zador, A.M. (2016). High-Throughput Mapping of Single-Neuron Projections by Sequencing of Barcoded RNA. Neuron 91, 975–987. 10.1016/j.neuron.2016.07.036.

Kishi, J.Y., Lapan, S.W., Beliveau, B.J., West, E.R., Zhu, A., Sasaki, H.M., Saka, S.K., Wang, Y., Cepko, C.L., and Yin, P. (2019). SABER amplifies FISH: enhanced multiplexed imaging of RNA and DNA in cells and tissues. Nat. Methods 16, 533–544. 10.1038/s41592-019-0404-0.

Li, Y., Walker, L.A., Zhao, Y., Edwards, E.M., Michki, N.S., Cheng, H.P.J., Ghazzi, M., Chen, T.Y., Chen, M., Roossien, D.H., et al. (2021). Bitbow Enables Highly Efficient Neuronal Lineage Tracing and Morphology Reconstruction in Single Drosophila Brains. Front. Neural Circuits 15, 732183. 10.3389/fncir.2021.732183.

Lichtman, J.W., and Denk, W. (2011). The Big and the Small: Challenges of Imaging the Brain’s Circuits. Science 334, 618–623. 10.1126/science.1209168.

Lillvis, J.L., Otsuna, H., Ding, X., Pisarev, I., Kawase, T., Colonell, J., Rokicki, K., Goina, C., Gao, R., Hu, A., et al. (2022). Rapid reconstruction of neural circuits using tissue expansion and light sheet microscopy. eLife 11, e81248. 10.7554/eLife.81248.

Lin, A., Yang, R., Dorkenwald, S., Matsliah, A., Sterling, A.R., Schlegel, P., Yu, S., McKellar, C.E., Costa, M., Eichler, K., et al. (2024). Network statistics of the whole-brain connectome of Drosophila. Nature 634, 153–165. 10.1038/s41586-024-07968-y.

Livet, J., Weissman, T.A., Kang, H., Draft, R.W., Lu, J., Bennis, R.A., Sanes, J.R., and Lichtman, J.W. (2007). Transgenic strategies for combinatorial expression of fluorescent proteins in the nervous system. Nature 450, 56–62. 10.1038/nature06293.

Matsliah, A., Yu, S., Kruk, K., Bland, D., Burke, A.T., Gager, J., Hebditch, J., Silverman, B., Willie, K.P., Willie, R., et al. (2024). Neuronal parts list and wiring diagram for a visual system. Nature 634, 166–180. 10.1038/s41586-024-07981-1.

McClure, C.D., Hassan, A., Aughey, G.N., Butt, K., Estacio-Gómez, A., Duggal, A., Ying Sia, C., Barber, A.F., and Southall, T.D. (2022). An auxin-inducible, GAL4-compatible, gene expression system for Drosophila. eLife 11, e67598. 10.7554/eLife.67598.

Motta, A., Berning, M., Boergens, K.M., Staffler, B., Beining, M., Loomba, S., Hennig, P., Wissler, H., and Helmstaedter, M. (2019). Dense connectomic reconstruction in layer 4 of the somatosensory cortex. Science 366, eaay3134. 10.1126/science.aay3134.

Owald, D., Fouquet, W., Schmidt, M., Wichmann, C., Mertel, S., Depner, H., Christiansen, F., Zube, C., Quentin, C., Körner, J., et al. (2010). A Syd-1 homologue regulates pre- and postsynaptic maturation in Drosophila. J. Cell Biol. 188, 565–579. 10.1083/jcb.200908055.

Park, S.Y., Sheridan, A., An, B., Jarvis, E., Lyudchik, J., Patton, W., Axup, J.Y., Chan, S.W., Damstra, H.G.J., Leible, D., et al. (2025). Combinatorial protein barcodes enable self-correcting neuron tracing with nanoscale molecular context. 2025.09.26.678648. 10.1101/2025.09.26.678648.

Santuy, A., Tomás-Roca, L., Rodríguez, J.-R., González-Soriano, J., Zhu, F., Qiu, Z., Grant, S.G.N., DeFelipe, J., and Merchan-Perez, A. (2020). Estimation of the number of synapses in the hippocampus and brain-wide by volume electron microscopy and genetic labeling. Sci. Rep. 10, 14014. 10.1038/s41598-020-70859-5.

Schlegel, P., Yin, Y., Bates, A.S., Dorkenwald, S., Eichler, K., Brooks, P., Han, D.S., Gkantia, M., dos Santos, M., Munnelly, E.J., et al. (2024). Whole-brain annotation and multi-connectome cell typing of Drosophila. Nature 634, 139–152. 10.1038/s41586-024-07686-5.

Schwanhäusser, B., Busse, D., Li, N., Dittmar, G., Schuchhardt, J., Wolf, J., Chen, W., and Selbach, M. (2011). Global quantification of mammalian gene expression control. Nature 473, 337–342. 10.1038/nature10098.

Shah, S., Takei, Y., Zhou, W., Lubeck, E., Yun, J., Eng, C.-H.L., Koulena, N., Cronin, C., Karp, C., Liaw, E.J., et al. (2018). Dynamics and Spatial Genomics of the Nascent Transcriptome by Intron seqFISH. Cell 174, 363–376.e16. 10.1016/j.cell.2018.05.035.

Shapson-Coe, A., Januszewski, M., Berger, D.R., Pope, A., Wu, Y., Blakely, T., Schalek, R.L., Li, P.H., Wang, S., Maitin-Shepard, J., et al. (2024). A petavoxel fragment of human cerebral cortex reconstructed at nanoscale resolution. Science 384, eadk4858. 10.1126/science.adk4858.

Shi, H., He, Y., Zhou, Y., Huang, J., Maher, K., Wang, B., Tang, Z., Luo, S., Tan, P., Wu, M., et al. (2023). Spatial atlas of the mouse central nervous system at molecular resolution. Nature 622, 552–561. 10.1038/s41586-023-06569-5.

Stark, W.M., Boocock, M.R., and Sherratt, D.J. (1992). Catalysis by site-specific recombinases. Trends Genet. TIG 8, 432–439.

Su, J.-H., Zheng, P., Kinrot, S.S., Bintu, B., and Zhuang, X. (2020). Genome-Scale Imaging of the 3D Organization and Transcriptional Activity of Chromatin. Cell 182, 1641–1659.e26. 10.1016/j.cell.2020.07.032.

Takei, Y., Zheng, S., Yun, J., Shah, S., Pierson, N., White, J., Schindler, S., Tischbirek, C.H., Yuan, G.-C., and Cai, L. (2021). Single-cell nuclear architecture across cell types in the mouse brain. Science 374, 586–594. 10.1126/science.abj1966.

Takei, Y., Yang, Y., White, J., Goronzy, I.N., Yun, J., Prasad, M., Ombelets, L.J., Schindler, S., Bhat, P., Guttman, M., et al. (2025). Spatial multi-omics reveals cell-type-specific nuclear compartments. Nature 641, 1037–1047. 10.1038/s41586-025-08838-x.

Tavakoli, M.R., Lyudchik, J., Januszewski, M., Vistunou, V., Agudelo Dueñas, N., Vorlaufer, J., Sommer, C., Kreuzinger, C., Oliveira, B., Cenameri, A., et al. (2025). Light-microscopy-based connectomic reconstruction of mammalian brain tissue. Nature 642, 398–410. 10.1038/s41586-025-08985-1.

Viswanathan, S., Williams, M.E., Bloss, E.B., Stasevich, T.J., Speer, C.M., Nern, A., Pfeiffer, B.D., Hooks, B.M., Li, W.-P., English, B.P., et al. (2015). High-performance probes for light and electron microscopy. Nat. Methods 12, 568–576. 10.1038/nmeth.3365.

White, J.G., Southgate, E., Thomson, J.N., and Brenner, S. (1986). The structure of the nervous system of the nematode Caenorhabditis elegans. Philos. Trans. R. Soc. Lond. B. Biol. Sci. 314, 1–340. 10.1098/rstb.1986.0056.

Winding, M., Pedigo, B.D., Barnes, C.L., Patsolic, H.G., Park, Y., Kazimiers, T., Fushiki, A., Andrade, I.V., Khandelwal, A., Valdes-Aleman, J., et al. (2023). The connectome of an insect brain. Science 379, eadd9330. 10.1126/science.add9330.

Xu, C.S., Hayworth, K.J., Lu, Z., Grob, P., Hassan, A.M., García-Cerdán, J.G., Niyogi, K.K., Nogales, E., Weinberg, R.J., and Hess, H.F. (2017). Enhanced FIB-SEM systems for large-volume 3D imaging. eLife 6, e25916. 10.7554/eLife.25916.

Yao, Z., van Velthoven, C.T.J., Kunst, M., Zhang, M., McMillen, D., Lee, C., Jung, W., Goldy, J., Abdelhak, A., Aitken, M., et al. (2023). A high-resolution transcriptomic and spatial atlas of cell types in the whole mouse brain. Nature 624, 317–332. 10.1038/s41586-023-06812-z.

Zhang, M., Pan, X., Jung, W., Halpern, A.R., Eichhorn, S.W., Lei, Z., Cohen, L., Smith, K.A., Tasic, B., Yao, Z., et al. (2023). Molecularly defined and spatially resolved cell atlas of the whole mouse brain. Nature 624, 343–354. 10.1038/s41586-023-06808-9.

Zheng, Z., Lauritzen, J.S., Perlman, E., Robinson, C.G., Nichols, M., Milkie, D., Torrens, O., Price, J., Fisher, C.B., Sharifi, N., et al. (2018). A Complete Electron Microscopy Volume of the Brain of Adult Drosophila melanogaster. Cell 174, 730–743.e22. 10.1016/j.cell.2018.06.019.

